# Prelimbic cortical excitatory overdrive and inhibitory underdrive accompany environmental suppression of food seeking

**DOI:** 10.1101/2025.05.21.655312

**Authors:** Kate Z. Peters, Zuzana Pedan, Romarua Agbude, Emily C. Woods, Oliver G. Steele, Nobuyoshi Suto, Scott B. Kinghorn, Olga Tsaponina, Eisuke Koya

## Abstract

Cues associated with food, such as fast-food advertising, can provoke food cravings and may lead to unhealthy overeating. Environmental enrichment (EE) that enhances cognitive and physical stimulation can reduce cue-evoked sucrose seeking in mice and recruitment of sucrose cue-reactive neurons or ‘neuronal ensembles’ in the prelimbic cortex (PL), which regulates appetitive behaviors. Hence, EE provides us with a behavioral model and neuronal targets to identify ‘anti-craving’ relevant mechanisms. Here, we investigated in the PL how EE modulated neuronal excitability and activity patterns in cue-reactive neuronal populations. Chemogenetic inhibition of cue-reactive neurons in PL blocked cue-evoked sucrose seeking, thereby confirming the function of these neurons in sucrose cue memory. EE boosted the baseline excitability of ‘originally’, or before EE exposure, cue-reactive, excitatory pyramidal cells in PL. Furthermore, their sucrose cue-specificity was lost – resulting in their persistent activation and non-cue selective activation or ‘excitatory overdrive’. Furthermore, EE reduced recruitment of cue-reactive, inhibitory interneurons reflecting ‘inhibitory underdrive’. Taken together, impaired neuronal food cue processing due to simultaneous prefrontal cortical excitatory ‘overdrive’ and inhibitory ‘underdrive’ likely underlies EE’s anti-craving action, thereby serving as potential neurophysiological targets to develop novel medications that help control food cravings.

## Introduction

External stimuli or ‘cues’, such as fast-food advertisements, associated with palatable foods trigger the retrieval of food memories and can provoke food cravings that promote unhealthy overeating [1,2]. Likewise in laboratory animals, food cues provoke motivated behaviors such as food seeking or ‘cue-evoked food seeking’ [3–5]. Therefore, to better understand how cues can provoke such reactions, considerable efforts have been made to reveal the corresponding brain mechanisms with the goal of intervening with the ‘pro-craving’ system. However, more needs to be established about how the brain ‘suppresses’ food cue reactivity. Such research is critical for understanding how the brain could harness its existing neuronal circuits to dampen cue reactivity and thus promote ‘anti-craving’ relevant effects.

Interestingly, cognitive and physical stimulation, including playing games and physical exercise reduces attentional bias towards food cues and subjective food cravings [6–8]. We and others have provided such stimulation to laboratory rodents by providing housing with environmental enrichment (EE). This enriched housing includes aspects such as larger cages with exercise wheels, tunnels and shelters, toys, and multiple nesting materials compared to standard housing (SH, **Figs. 1a, b)** [5,9–11]. In line with humans, EE suppresses cue-evoked sucrose seeking in both mice and rats [5,11]. Furthermore, EE attenuated recruitment of sparse sets of cue-reactive neurons or ‘neuronal ensembles’ expressing the neuronal activity marker ‘Fos’ in the prelimbic cortex (PL) [5,12], a region of the medial prefrontal cortex (mPFC), that controls motivated actions [13–15]. Therefore, EE provides us with a useful behavioral procedure and prefrontal neuronal targets to identify anti-craving effects.

**Figure 1.**
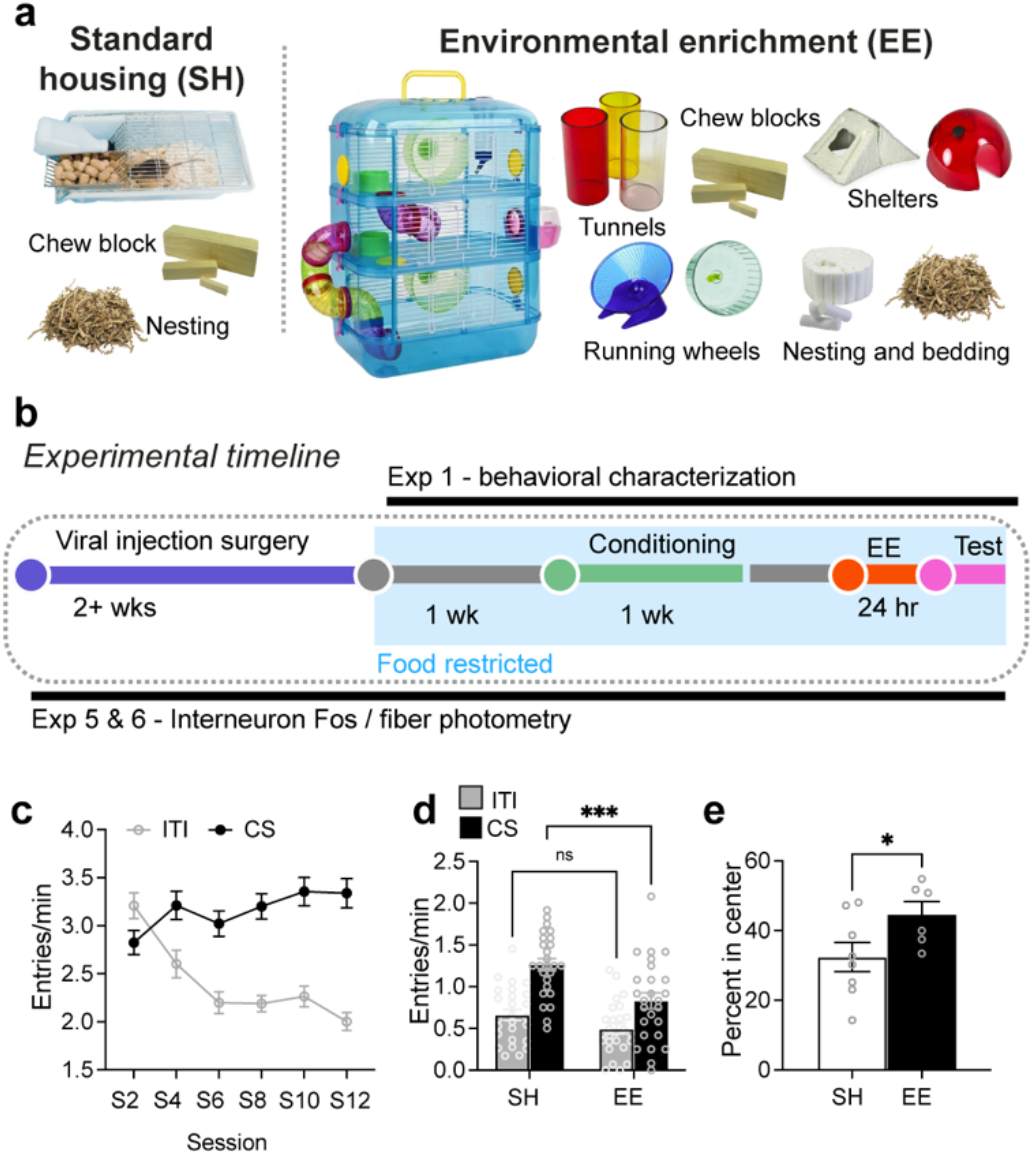
Experimental timelines and housing conditions. **a** images of housing with environmental enrichment (EE) and control standard housing (SH), **b** Timeline. **c** Head entry rates into the magazine during the conditioned stimulus (CS, cue) and intertrial interval (ITI, no cue) periods during acquisition for all mice. **d** Head entry rates at test session (n=26/group). **e** percent time spent in center of open field (n=6-8/group). *p < 0.05 and ***p< 0.001. All data are expressed as mean ± SEM.

We previously revealed that the modulation of cue-evoked food seeking is controlled via alterations in mPFC neuronal ensemble recruitment and excitability [16,17]. Furthermore, rapid changes in the *in vivo* activity patterns of mPFC neurons interpret the changes in meaning of reward-associated cues [18–20]. Here we hypothesized that EE promoted the suppression of cue-evoked food seeking through modulating the excitability- and activity-related properties of PL neurons. We thus investigated these properties in the PL in sucrose conditioned mice that learned a ‘cue-food’ association and were exposed to EE.

## Materials and Methods

For detailed information see supplementary methods. Wild-type (with GCaMP6f expression (only females) or without (males and females), *FosTRAP2* expressing Cre-dependent hM4Di-mCherry (male and female), *FosTRAP2:Ai14* (only females) were used. *FosTRAP2* or *FosTRAP2:Ai14* mice received 4-OHT (50 mg/kg, i.p.) to TRAP with hM4Di-mCherry or tdTomato, respectively, 3 hours following initiation of a sucrose seeking test. To activate hM4Di, *FosTRAP2* mice received clozapine (0.1 mg/kg, i.p.). Mice underwent sucrose conditioning, testing for sucrose seeking in Med Associated conditioning chambers, EE exposure, and locomotor activity testing as described previously [5,21].

Histological brain sections were prepared and visualized similar to our previous studies [4,17,21]. PL-containing sections were imaged and analyzed for immunofluorescence (Fos) and/or native fluorescence (mCherry, mRuby, tdTomato, GCaMP6f) using iVision software.

Brain slice electrophysiology was performed similar to as described previously [4,17,21]. Data was analyzed using Easy Electrophysiology software.

A TDT RZ10x fiber photometry system and Synapse software were used for GCaMP fiber photometry data collection. For all experiments, data was analyzed using ANOVAs followed by post-hoc testing or pair-wise comparisons using Prism software (RRID:SCR_002798; GraphPad 10 Software). All experiments were conducted in accordance with the UK Animals (Scientific Procedures) Act 1986 (ASPA, amended 2012) and received ethical approval from the University of Sussex Animal Welfare and Ethics Review Board (AWERB).

## Results

### Experiment 1 – EE reduces cue-evoked food seeking, but not general locomotor activity

#### Pavlovian Conditioning

We conditioned mice to associate an auditory cue which served as a conditioned stimulus (CS) with 10% sucrose solution delivery across 12 sessions **(Fig. 1c**). There was a significant Session x CS interaction (F_11, 578_ = 25.5, p<0.001), and main effects of session (F_11,605_ = 5.7, p<0.001) and CS (F_1,55_ = 144.0, p<0.001). Thus, across training sessions mice acquired a cue-food association, i.e. a food cue memory.

#### Cue-evoked sucrose seeking

We assessed effects of EE on cue-evoked sucrose seeking with a Test session under extinction conditions. There was a significant interaction of EE x CS (F_1,50_ = 6.371, p<0.05), demonstrating that EE selectively attenuated cue-evoked sucrose seeking (**Fig. 1d**). We also observed a significant main effect of CS (F_1,50_ = 10.42, p<0.01) and EE (F_1,50_ = 79, p<0.001). Post-hoc tests showed EE selectively reduced CS responding and had no effect on inter-trial interval (ITI or no cue period) responding, indicating that the reduction in sucrose seeking was not due to a general suppression of locomotor activity.

#### Locomotor activity tests

A subset of mice underwent a locomotor activity test. There was no significant difference in total distance travelled between SH and EE mice (t_(12)_=0.376, p=0.713; **Fig. S1a**). However, mice in the EE condition spent a significantly higher percentage of their time in the center of the arena compared to SH mice (U=8, p<0.05; **Fig. 1e**) suggesting a reduction in anxiety-like behavior within the open field test [22].

### Experiment 2: Silencing a cue-reactive PL ensemble inhibits sucrose seeking

Sparse sets of cue-reactive neurons or ‘neuronal ensembles’ in mPFC establish appetitive, ‘cue-reward’ memories and modulate cue-evoked food seeking [17,23]. Since EE attenuates cue-evoked sucrose seeking, a behavior that requires establishing cue-food associations, we investigated whether EE exerts its effects by modulating the excitability and recruitment properties of the originally, i.e. before EE exposure, cue-reactive PL ensemble. We first examined whether the PL contained a neuronal ensemble that mediated cue-evoked sucrose seeking and thus established a food cue memory in *FosTRAP2* mice [24]. Here, PL neurons were tagged with the inhibitory DREADD ‘hM4Di’ [25] using 4-hydroxy-tamoxifen (4-TM) during cue-evoked sucrose seeking.

#### Demonstration of 4-TM-induced tagging of hM4di-mCherry following conditioning

To demonstrate stimuli- and 4-TM dependence of tagging in *FosTRAP2* mice, we assessed levels of mCherry+ cells in PL from 3 groups of *FosTRAP2* mice that received PL injections of AAV-DIO-hM4Di-mCherry (**Fig. 2a**). Mice in these groups received 4-TM injections following a conditioning session (4-TM+/S) or in the home cage (4-TM+/HC) or those that did not receive 4-TM injections in the home cage (4-TM–/HC).

**Figure 2.**
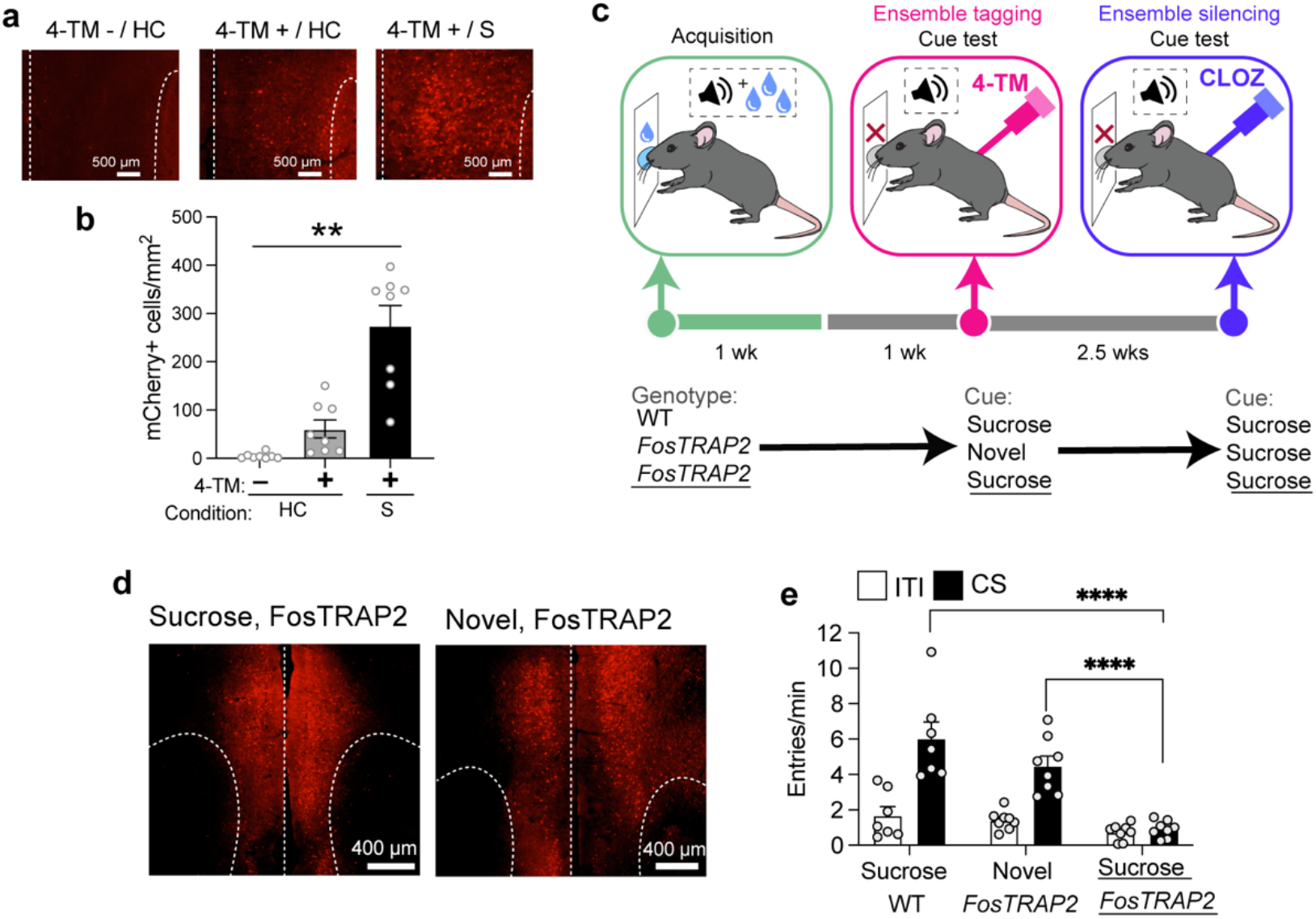
The 4-hydroxy-tamoxifen (4-TM) dependency of tagging with hM4Di-mCherry and silencing (prelimbic) PL ensembles tagged with hM4di-mCherry disrupts cue-evoked sucrose seeking. **a** Representative PL images and **b** levels of hM4Di-mCherry expression from mice not injected with 4-TM in home cage (4-TM-/HC; n = 8); injected with 4-TM in home cage (4-TM+/HC; n = 8) and following a conditioning session (4-TM+/S); n = 8). **c** Experimental timeline for tagging PL ensembles with hM4Di-mCherry during cue-evoked sucrose seeking and ensemble silencing on test day. **d** Representative images of PL tagged with hM4Di-mCherry following sucrose cue and novelty exposure. **e** Disruption of cue-evoked sucrose seeking following clozapine in *FosTRAP2* mice tagged with hM4Di-mCherry following sucrose cue exposure (Sucrose *FosTRAP2*, n = 8); but not following novelty (Novel *FosTRAP2*, n = 8); and WT control mice that underwent sucrose cue exposure (Sucrose WT, n = 7). *p < 0.05, **p<0.01 and ****p< 0.0001. All data are expressed as mean ± SEM.

A 1-way ANOVA revealed significant differences between the groups in mCherry counts (F_2, 21_ = 28.91, p<0.0001; **Fig. 2b**). Post-hoc testing revealed significantly higher levels of mCherry+ cells in the 4-TM+/S compared to the control group (4-TM+/HC and 4-TM–/HC) and higher levels of mCherry+ cells in the 4-TM+/HC compared to 4-TM-/HC group. Therefore, tagging was 4-TM-dependent and most robustly occurred following presentation of appetitive stimuli that enhances PL Fos expression [26].

#### hM4Di-silencing of PL ensembles

We examined whether the ‘originally’ cue-reactive PL ensembles activated during sucrose cue exposure would play a causal role in cue-evoked sucrose seeking (**Fig. 2c)**. We injected AAV-DIO-hM4Di-mCherry in the PL in 2 groups of *FosTRAP2* and 1 group of wild-type mice. One week following conditioning, all mice were injected with 4-TM following a cue-evoked sucrose seeking test for tagging (or no tagging in WT mice) cue- and novelty-activated ensembles (**Fig. 2d**) with hM4Di in *FosTRAP2* mice. The novelty-tagged control was included as novel stimuli robustly activate neuronal ensembles that are distinct from those activated by reward-associated cues [27,28].

Approximately 2 weeks following tagging on test day, all mice received behaviorally [17] and neuronally [29] inert, subthreshold injections of the DREADD agonist clozapine [30] (0.1 mg/kg, i.p., **Fig. S2**).

We next measured the effects of ensemble silencing on cue-evoked sucrose seeking. A 2-Way ANOVA revealed a significant Group x CS interaction (F_2, 20_ = 25.06, p < 0.0001). Post hoc analysis revealed that sucrose cue-tagged *FosTRAP2* mice did not exhibit cue-evoked sucrose seeking (**Fig. 2e)**, whereas novelty-tagged *FosTRAP2* mice and sucrose-tagged WT mice did. These data indicate that the PL contains a neuronal ensemble that mediates cue-evoked sucrose seeking.

### Experiment 3– EE upregulates baseline firing capacity in cue-reactive PL pyramidal cell ensembles

We previously revealed that the majority (>97%) of cue-reactive, *Fos*-expressing prefrontal ensembles were excitatory, pyramidal cells [4]. We examined whether the identity of cue-tagged PL neurons were pyramidal cells and whether EE altered the excitability of PL neuronal ensembles that mediate cue-evoked sucrose seeking in *FosTRAP2:Ai14* mice. One week following the final conditioning session, these mice underwent a cue-evoked sucrose seeking test and their activated neurons were tagged with the red fluorescent reporter ‘tdTomato’ using 4-TM. Subsequently, >2 weeks later mice were either exposed to 1d EE or remained in their standard housing (SH) and we examined excitability alterations in pyramidal cell ensemble (tdTomato+) and non-ensemble (tdTomato–) neurons in PL (**Fig. 3a, b**).

**Figure 3.**
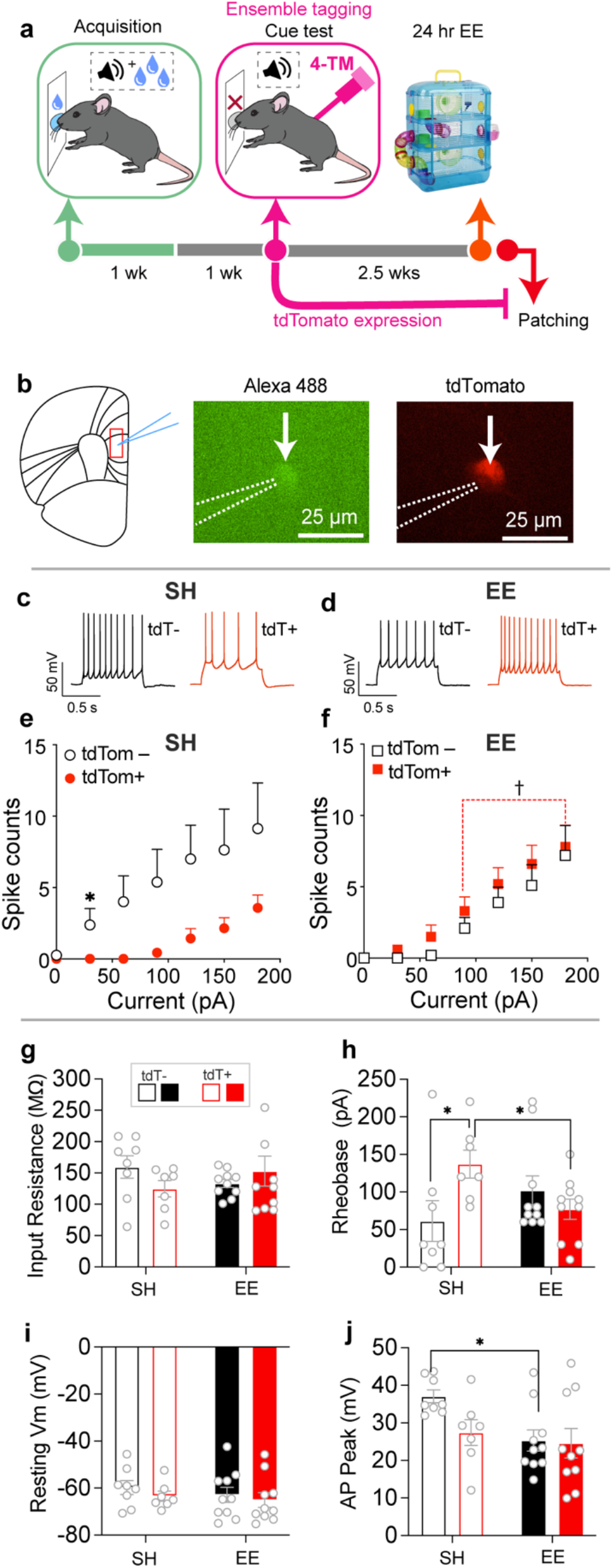
EE upregulated the excitability of ensemble, but not non-ensemble neurons in PL. **a** Experimental timeline. **b** Representative image of tdTomato+ cell recording (indicated by white arrow) in PL. Respective representative current traces from **c** tdTomato– (tdT–) and **d** tdTomato+ (tdT+) cells. The firing capacity **e** tdTomato– (tdTom–) and **f** tdTomato+ (tdTom+) cells. g**-j** Membrane properties of tdTomato– (tdTom–) and tdTomato+ (tdTom+) cells (EE: n=10/4, tdTomato+ and 9/5, tdTomato; SH: n=7/4, tdTomato+ and n=8/3, tdTomato–). *n* = total number of cells/total number of mice. ^*^p<0.05; ^†^p<0.05 tdTomato+ cells in EE vs SH. All data are expressed as mean ± SEM.

Consistent with our previous studies [4,17], we found that all cells identified via their tdTomato expression (17 out of 17 tdTomato+ cells) and non-expression (17 out of 17 tdTomato– cells) exhibited firing patterns reminiscent of pyramidal cells, similar to that shown in **Figures 3c,d**. Hence, our cue-tagged ensemble neurons in *FosTRAP2* mice are primarily pyramidal cells. We next examined firing capacity changes following current injections in these neuronal populations. A 3-way ANOVA revealed a significant effect of EE x Current x tdTomato (F_18, 558_ = 2.002; p<0.01; **Fig. 3e,f**). To reveal the source of this interaction, we examined the firing capacity of tdTomato+ and – populations within EE and SH conditions using a 2-way ANOVA. Ensemble neurons exhibited downregulated firing capacity compared to non-ensemble neurons in the SH, but not EE condition (Current x tdTomato; SH: F_18,234_ = 2.388, p<0.01; EE: F_18,324_ = 0.3129; p = 0.9972; **Fig. 3e,f**).

We also examined these properties within tdTomato+ and – populations across EE and SH conditions. EE upregulated firing capacity in ensemble, but not non-ensemble neurons (EE x Current; ensemble neurons: *F*_18,270_ = 4.063; *p*<0.0001, **Fig. S3**; non-ensemble neurons: *F*_18,288_= 0.6941; *p* = 0.8166, **Fig. S3**). We next examined the underlying intrinsic adaptations that may contribute to the increased firing capacity of tdTomato+ neurons in the EE group for the parameters in **Figures 3g-j and Table S1**. The observed upregulation was attributable to a reduction in the depolarization threshold, indicated by decreased rheobase (EE x tdTomato; *F*_1,31_ = 6.351; p <0.05; **Fig. 3h**) which may reflect increases in the number and/or function of voltage-gated Na^+^ channels [31]. Post-hoc testing revealed a significantly enhanced rheobase in tdTomato+ compared to tdTomato– cells in the SH condition and reduced rheobase in tdTomato+ cells in the EE compared to SH condition. These data indicate that EE upregulated the firing capacity of tdTomato+, but not tdTomato– cells, and tdTomato+ cells exhibited downregulated firing capacity compared to tdTomato– cells in the SH condition.

For other membrane parameters, EE did not regulate other active and passive membrane properties in ensemble neurons indicated by their lack of significant EE x tdTomato interactions (**Fig. 3g, i, j** and **Table S1**). However, there was a main effect of EE for AP peak (*F*_1,31_ = 5.131; *p*<0.05, **Fig. 3j**) and AP half-width (*F*_1,31_ = 4.673; *p*<0.05, **Table S1**). Post-hoc testing revealed a significantly lower peak and half-width in tdTomato– cells in the EE compared to SH condition.

### Experiment 4– EE induced persistent activation of cue-reactive PL ensemble

Since we revealed a PL ensemble that mediated cue-evoked sucrose seeking and a pyramidal cell ensemble that exhibited upregulated excitability following EE, we investigated whether EE modulated the recruitment of this ensemble during a subsequent cue-evoked sucrose seeking test in *FosTRAP2:Ai14* mice (**Fig. 4a**). One week following the final conditioning session, *FosTRAP2:Ai14* mice underwent a tagging session during cue-evoked sucrose seeking. Their activated neurons were tagged with tdTomato following 4-TM injections (**Fig. 4a, b)**. Subsequently, >2 weeks later, mice were either exposed to 1d EE or remained in standard housing (SH) and then underwent a sucrose seeking test. A 2-way ANOVA revealed a significant interaction effect of EE x CS, (F_1, 19_ = 5.57, p < 0.05) and a main effect of CS (F_1, 19_ = 13.69, p < 0.01) and EE (F_1, 19_ = 18.20, p < 0.01; **Fig. 4c**), indicating that EE selectively reduced cue-evoked sucrose seeking.

**Figure 4.**
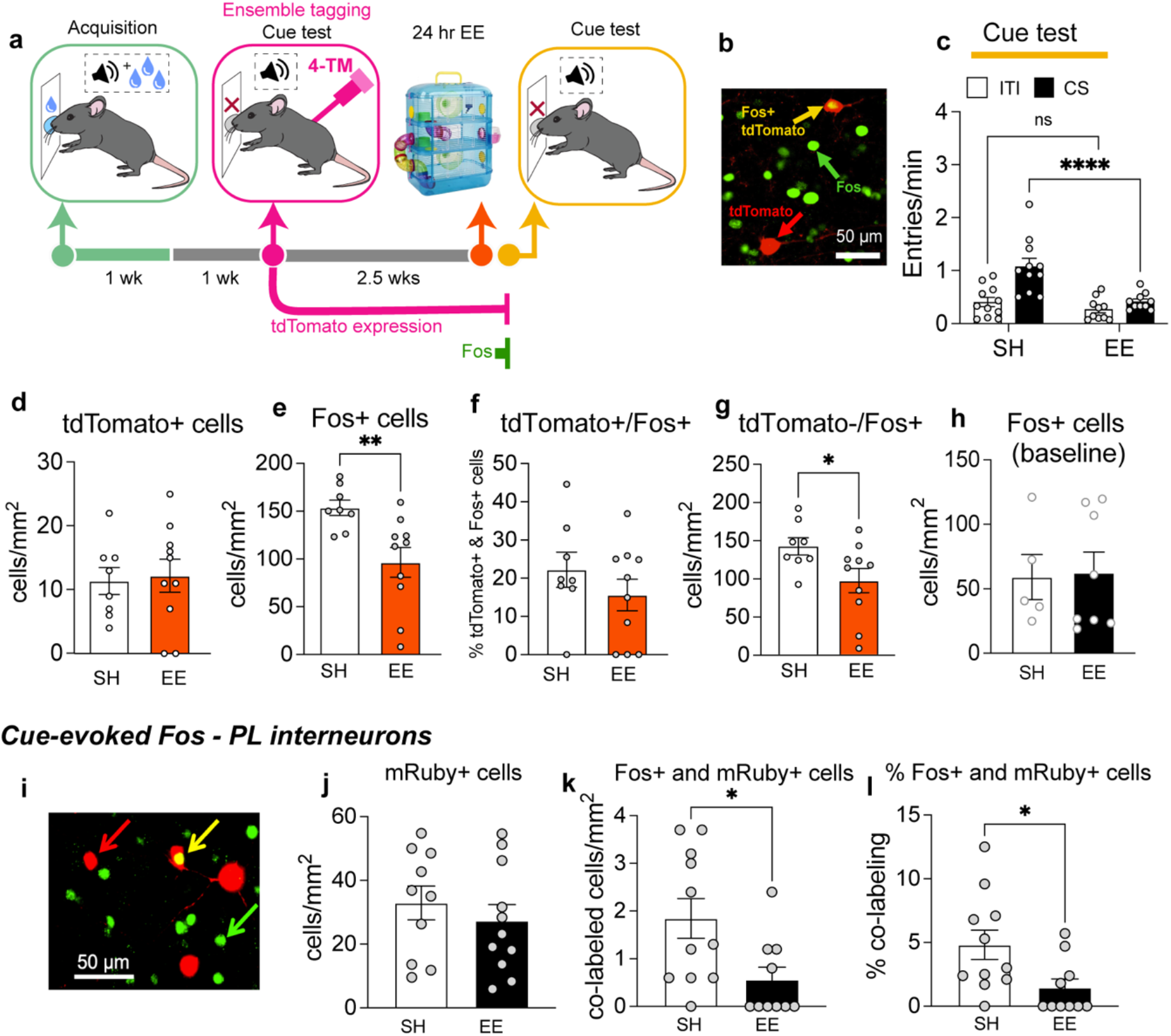
EE does not modulate the recruitment patterns of neuronal ensembles tagged with tdTomato during cue-evoked sucrose seeking in PL but reduces Fos in tdTomato– and mRuby+ cells. **a** Experimental timeline of tagging and cue testing. **b** Representative image of tdTomato+ (red arrow), Fos+ (green arrow), and tdTomato+/Fos+ (yellow arrow) cells. **c** EE attenuates cue evoked sucrose seeking during Cue test 2 (n=11/group). **d** Similar levels of tdTomato-tagging in mice from EE and SH conditions. **e** EE generally reduces Fos expression, **f** but not levels of tdTomato+/Fos+ expression. **g** EE reduces levels of tdTomato–/Fos+ expression (e-h; n=8-10/group). **h** EE does not modulate baseline Fos prior to cue exposure. EE attenuated cue-evoked Fos in interneurons. **i** Representative image of mRuby+ (red arrow), Fos+ (green arrow), and mRuby+/Fos+ (yellow arrow) cells. **j** Similar levels of mRuby expression across EE and SH conditions. **k** EE reduces levels of mRuby–/Fos+ cells and **l** only a minority of mRuby+ cells express Fos. (i-k; n=10-11/group). **p<0.01 and *p<0.05, All data are expressed as mean ± SEM.

Following this testing, we examined levels of tdTomato-tagging and Fos expression in PL. We found similar tdTomato-tagging levels from mice in EE and SH conditions (*t*_(16)_ = 0.24, *p* = 0.81; **Fig. 4c,d)**. EE significantly reduced cue-evoked Fos (*t*_(16)_ = 3.02, *p* < 0.01, **Fig. 4e**), similar to our previous study [5]. We next determined whether cue-tagged, ensemble neurons exhibited differential reactivation patterns following EE. We assessed the levels of tdTomato+ cells that co-expressed Fos (tdTomato+/Fos+ cells). Surprisingly, EE did not modulate the reactivation patterns of tdTomato+/Fos+ ensemble neurons (*t*_(15)_ = 0.32, *p* = 0.75; **Fig. 4f)**. However, EE reduced levels of tdTomato–/Fos+ cells and thus recruitment of non-tagged ensembles (*t*_(16)_ = 2.20, *p* < 0.05, **Fig. 4g**).

We next determined in a separate cohort of sucrose conditioned wild-type mice whether this decrease was due to a potential baseline (prior to cue exposure) Fos decrease in PL following EE access, one week following the final conditioning session. We observed no significant differences in baseline Fos (*t*_(11)_ = 0.136, *p* = 0.895; **Fig. 4h)**. Therefore, EE’s dampening effects on levels of Fos+ and tdTomato–/Fos+ cells (**Fig. 4e,g**) and in our previous study [5] is more associated with reductions in sucrose seeking, rather than attenuating baseline PL Fos expression.

### Experiment 5 – EE decreases cue-evoked Fos in interneurons

Since EE reduced PL Fos expression and induced ensemble reactivation, we next investigated whether this was linked to reduced PL inhibition. To this end, mice were injected with AAV-mDlx-mRuby [32] in the PL to express the red fluorophore ‘mRuby’ in inhibitory, GABAergic interneurons (**Fig. 4i**) and we assessed Fos expression in mRuby-expressing interneurons. Similar mRuby expression was observed in mice from EE and SH conditions (*t*_(19)_ = 0.761, *p* = 0.456; (**Fig 4j**). EE significantly reduced levels of Fos+ and mRuby+ cells (*t*_(19)_ = 2.551, p=0.0195; (**Fig. 4k**) and percentage of mRuby+. Cells that expressed Fos (*U =* 21.5, p=0.015; **Fig. 4l**). Only a minority of mRuby+ cells expressed Fos; on average 2.06% following EE and 5.40% following SH.

## Experiment 6 – EE induces loss of cue specificity and general enhancements of PL activity *in vivo*

### GCaMP activity during Pavlovian conditioning

To reveal how EE rapidly modulates neuronal activity during suppression of sucrose seeking, we recorded PL calcium activity *in vivo* in pyramidal cells using fiber photometry [33] (**Fig. 5a, b**). we quantified activity peaks (transients) a median absolute deviation (M.A.D) threshold method [34] (**Fig. 5c**).

**Figure 5.**
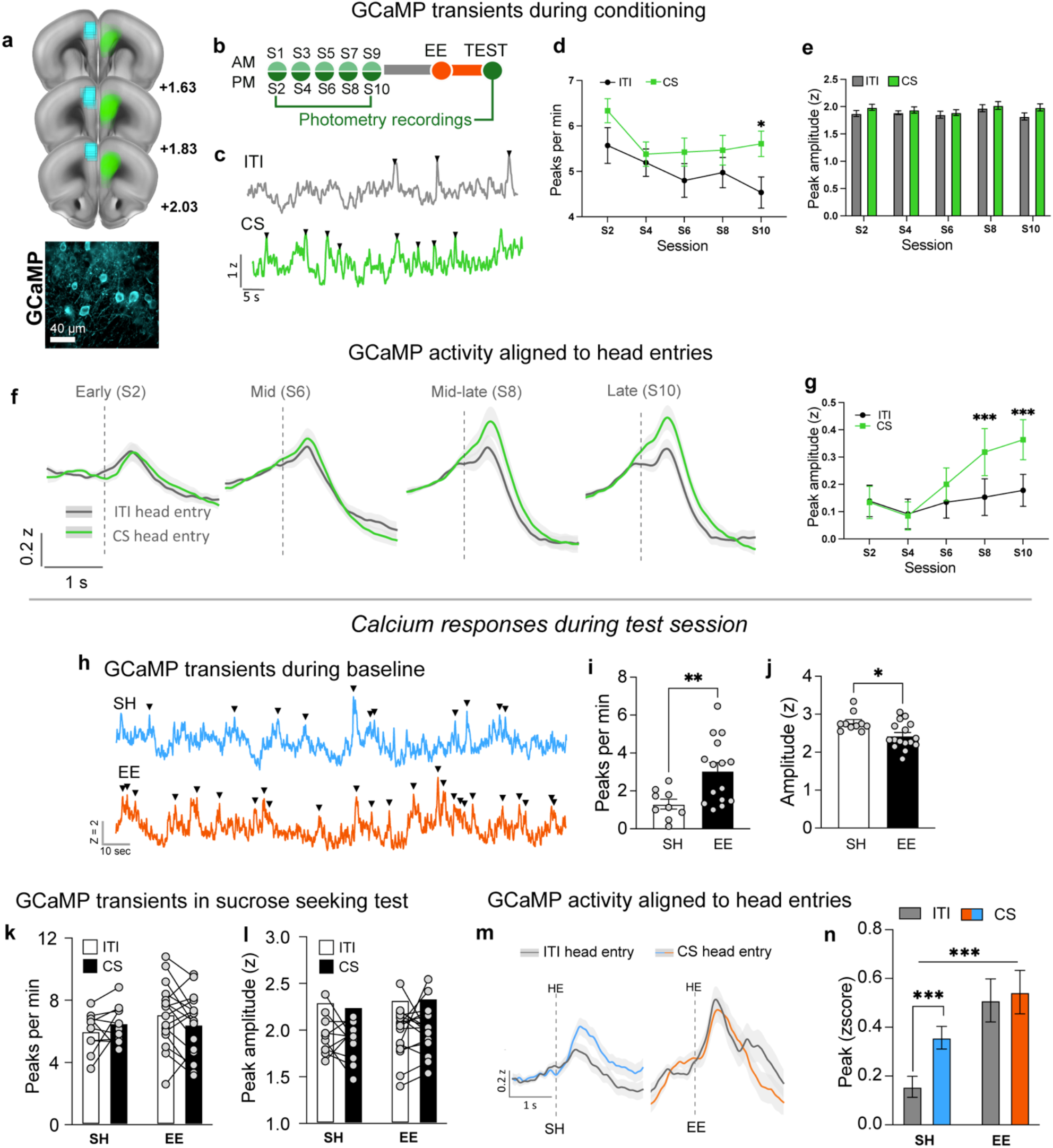
Acquisition of sucrose conditioning is associated with cue-specific *in vivo* activity of PL pyramidal cells (n=26) and EE induced loss of cue-specific and generally elevated PL activity *in vivo* during sucrose seeking (n=10-16/group). **a** Viral expression patterns and fiber placement for PL GCaMP recordings. **b** Timeline **c** Representative GCaMP transients during conditioning, activity during ITI (grey) and CS (green), black triangles indicated identified peaks. **d** Frequency of transient calcium events (peaks per minute) during ITI (black) and CS (green) across conditioning sessions. By session 10 (S10) the frequency of calcium transients is significantly higher during the CS period compared to ITIs. **e** Mean peak amplitude of transients during ITIs (grey) and CSs (green) across conditioning sessions. **f A**veraged traces for head entries in CS and ITI periods across conditioning. Activity was time locked to head entry onset and events within 2 seconds of head entry were averaged. Average traces are shown for early (S2), mid (S6), mid—late (S8) and late (S10) conditioning stages. Grey traces show neural responses during ITI and green during CS. **g** Mean peak amplitudes during conditioning sessions. Mean peak amplitude – highest averaged activity within a 2 second window – is expressed as z-score is shown across conditioning. By session 8 averaged peak signals are significantly higher during CS than during ITI. **h** Representative GCaMP transients during 15-minute baseline period, standard housed (SH – blue) and environmentally enriched (EE – orange). **i** Frequency of transient calcium events (peaks per minute) during baseline. EE (black bars) show significantly higher frequency of events that SH (white bars). **j** Mean peak amplitude of transients during baseline period. EE (black bar) reduced the mean amplitude of peaks compared with SH (white bar). **k** Frequency of transient calcium events (peaks per minute) during ITI (white bars) and CS (black bars) in SH (left) and EE (right) mice. **l** Mean peak amplitude of transients during ITI (white bars) and CSs (black bars). **m** Entry-related averaged traces for ITI (black lines) and CS (colored lines) periods during sucrose seeking test session, in SH (blue line) and EE (orange line). Activity was time locked to head entry onset and events within 2 seconds of head entry were averaged. **n** Entry-related mean peak amplitude – highest averaged activity within a 2 second window – is expressed as z-score is shown for ITI (grey bars) and CS (colored bars) in SH (blue) and EE (orange) mice. *p<0.05, **p<0.01, ***p<0.001. All data are expressed as mean ± SEM.

During conditioning, the frequency of transient events (peaks per minute) was significantly elevated in the CS compared to ITI periods (main effect of cue F_1,25_ = 19.69, p<0.001; **Fig. 5d**). *A priori* analyses using a t-test revealed significantly more peaks per min during CS versus ITI periods in the final conditioning session (Session 10, *t*_(23)_ = 3.047, p<0.05). We quantified the amplitude of all detected peaks and found a main effect of cue (CS vs ITI), F_1,25_ = 7.317, p<0.05, but no significant effect of session (F_3.26, 81.4_ = 1.474, p = 0.225; **Fig. 5e**). Hence, the acquisition of sucrose conditioning is associated with altered encoding patterns in PL pyramidal cells, such that cue presentations enhance the frequency, but not amplitude of calcium activity events *in vivo*.

Since cue-evoked sucrose seeking is associated with increased head entries during the CS compared to ITI periods, we investigated whether calcium activity was different during these periods. Averaged traces time-locked to head entries in CS and ITI periods for all mice and EE and SH groups separately across conditioning are shown in **Figures 5f and S4, respectively**. We split conditioning into early, mid, mid-late and late conditioning. We found a significant main effect of cue (CS vs ITI) F_1,24_ = 23.08, p <0.001), a significant main effect of session (early, mid, mid-late and late) F_3,72_ = 5.30, p<0.01) and a significant Cue x Session interaction (F_2.9, 67.51_ = 5.93, p < 0.01; **Fig. 5g**). Post-hoc testing revealed significantly elevated activity levels during the CS compared to ITI period emerging towards the end of conditioning (mid-late Session 8) ITI vs CS (*t*_(23)_ = 4.678, p<0.001) and at late conditioning (Session 10) *t*_(25)_ = 4.828, p<0.001). Overall, the acquisition of sucrose conditioning is associated with the cue-specific *in vivo* activity of PL pyramidal cells.

### GCaMP activity during sucrose seeking test

#### Baseline

We included a 15-minute baseline period to assess whether EE modulates baseline GCaMP activity prior to testing (**Fig. 5h-j)**. During baseline, EE mice had significantly more calcium events per minute than SH mice (*t*_(22)_ = 2.892, p<0.01 **(Fig. 5i)**. However, these peak transients were significantly lower in amplitude in EE mice vs SH mice, *t*_(24)_ = 2.755, p<0.05 **(Fig. 5j)**.

#### Cue-evoked sucrose seeking test

As with conditioning sessions and baseline, we used the same peak detection method to quantify peak frequency and amplitude specifically during ITI and CS periods. Two-way ANOVAs revealed no significant main effect of cue (ITI vs CS) on the rate (F_1,24_ = 0.044, p = 0.836) or amplitude (F_1,24_ = 0.232, p = 0.635) of calcium events. Calcium responses during the test session do not differentially respond to the CS or ITI period. During the test session there were also no significant main effect of housing between SH and EE groups in the frequency (F_1,24_ = 0.591, p = 0.449; **Fig. 5k** or in the amplitude of these events (F_1,24_ = 1.10, p = 0.304; **Fig. 5l**).

As we did not see significant differences in calcium activity during CS or ITI time periods, we next investigated whether the recorded neurons respond differently during head entries made in the CS and ITI periods (averaging all head entry trials). We compared all head entry related trials in ITI vs CS and in SH vs EE conditions. Averaged traces during ITI and CS in SH and EE mice are shown in **Figure 5m**. We found a generalized increase in head entry evoked activity in the EE condition (F_1,560_ = 18.04, p<0.001; (**Fig. 5n)**. Regardless of whether head entries occurred during CS or ITI those in the EE group were systematically elevated. Similar to the final conditioning session, *a priori* analysis revealed in the SH condition there was a significantly higher CS compared to ITI, entry-related calcium activity (*t*_(560)_ = 2.923, p<0.001, **(Fig. 5n)**. Conversely, in the EE condition this neural discrimination between CS and ITI entries on food cue specificity was lost as *a priori* analysis did not reveal differences between head entry evoked peaks in CS and ITI periods (*t*_(560)_ = 2.091, p>0.05).

## Discussion

Here we investigated how EE, an experience that suppresses food seeking, modulates PL excitability and activity-related properties and we revealed changes related to excitatory overdrive and inhibitory underdrive. First, chemogenetic inhibition of PL neurons that were ‘originally’ cue-reactive, i.e. before EE exposure, blocked cue-evoked sucrose seeking, thereby confirming their functional role in food cue memory. Next, related to excitatory overdrive, EE upregulated the baseline excitability of an originally (before EE) cue-reactive, pyramidal cell ensemble that established a food cue memory. Following cue-evoked sucrose seeking, EE still induced reactivation of these ensembles. Additionally, EE induced a loss of cue specificity and general elevation of PL pyramidal cell activity *in vivo* during sucrose seeking. Finally, related to inhibitory underdrive, EE reduced recruitment of inhibitory interneurons, suggesting reduced local inhibition that permits pyramidal cell overdrive. Since the general suppression of PL activity is known to reduce reward seeking [13], our findings provide deeper insight into how both excitatory and inhibitory PL mechanisms accompany how EE protects against food cue reactivity. Together, we reveal a mechanism that is relevant to ‘anti-food seeking’ consisting of PL pyramidal cell overdrive and interneuron underdrive, where neurons engaged by food cues and EE are modified. We discuss below the implications of the EE-induced, neurophysiology- and activity-related alterations and their impact on cue-evoked sucrose seeking.

### EE-induces excitatory overdrive of cue-reactive PL ensemble

#### Enhanced baseline excitability of cue-reactive ensemble

Interestingly, EE enhanced the baseline (prior to cue exposure) excitability of pyramidal cells that belonged to an ‘originally’ (before EE) cue-reactive PL ensemble that mediated cue-evoked sucrose seeking. This demonstrates that EE modified the properties of pyramidal cells that established a learned food cue memory, as prefrontal ensembles are primarily composed of pyramidal cells [4,17,35]. This enhanced excitability might reflect a homeostatic, compensatory mechanism to decreased excitatory inputs to these ensemble neurons following EE, as we previously found decreased excitatory synaptic strength onto Fos-expressing, neuronal ensembles [36,37]. Another possibility is that the enhanced ensemble excitability results from forming new learned associations between the physical attributes of the EE cage (e.g. exercise wheel) and the rewarding EE experience [38]. In support, we previously observed enhanced prefrontal ensemble excitability following the initial, but not final, sucrose conditioning session as mice were learning about sucrose and the cues predicting their availability [17]. Thus, from this perspective EE may take advantage of the existing cue-food ensemble and modify its excitability to encode a new cue-reward association.

#### Reactivation of cue-reactive ensemble during sucrose seeking

Contrary to our expectation, EE did not modulate reactivation of the originally, cue-reactive neuronal ensembles that formed a sucrose cue memory, as cue-tagged neurons were still reactivated in *FosTRAP2:Ai14* mice. This result was surprising given that we previously saw reduced likelihood of reactivation of the ‘original’ mPFC neuronal ensembles that were persistently activated in conditioning as mice suppressed sucrose seeking during extinction learning [16] and the suppression of responding to conditioned cues is associated with reduced reactivation of ensemble neurons [39].

Although a similar number of ensemble neurons were reactivated across EE and standard housing conditions, EE may have recruited a ‘distinct’ set of neurons within the originally cue-reactive (tdTomato+) neurons compared to standard housing conditions. These distinctly recruited neurons may exert top-down control to brain structures that regulate the expression of conditioned responses, such as the basolateral amygdala (BLA) [40]. Thus, future studies need to investigate these possibilities by labelling specific projections of these cue-reactive neurons using retrograde tracers. Finally, the enhanced ensemble excitability following EE may render these neurons to become more readily activated and express Fos. We argue below that this overactivity in cue-reactive neurons might impair cue impair cue-specific responses during sucrose seeking.

#### PL activity *in vivo* prior to and during sucrose seeking and links to enhanced ensemble excitability

Fiber photometry combined with genetically encoded calcium sensors (GCaMP) measures bulk calcium activity *in vivo* from defined cell populations. Here, prior to cue exposure, EE mice exhibited increased mean frequency, but decreased mean amplitude, of calcium transients from PL pyramidal cells. An explanation for the former may be due to increased detection of activity events from increased pyramidal cell ensemble excitability and increased frequency of excitatory inputs. On the other hand, the latter may result from weaker excitatory inputs to PL from afferent structures including the BLA, whose activity is reduced during the suppression of cue-induced reward seeking [41], i.e. EE increases the frequency of baseline excitatory input events, while weakening each event. Furthermore, since action potential amplitude and half-width influences calcium influx [42], the reduced calcium transient amplitude may result from EE-induced decrease in action potential amplitude and half-width in tdTomato– neurons. Of note, unlike baseline excitability and Fos parameters, baseline calcium signals were obtained in the training chamber. Thus, future studies should also measure baseline calcium signals under less behaviorally arousing conditions in the home cage.

During sucrose seeking, we observed that EE impaired the cue specificity of cue-reactive, PL pyramidal cells and generally enhanced their activity *in vivo*. This lack of difference in cue-related activity resembled what was observed in early conditioning sessions, prior to establishment of a robust cue-food association. Since alterations in excitability correlate with alterations in bulk calcium signals [43], the increased peak responses may arise from the enhanced excitability of pyramidal cell ensembles following EE, combined with increased recruitment of excitatory afferents to PL such as ventral hippocampus and BLA neurons. Additionally, the decreased action potential amplitude in in tdTomato– neurons and decreased recruitment of Fos+ and tdTomato– neurons suggest a dominant contribution of calcium signals from pyramidal cell ensembles. The increase in peak responses are thought to indicate an increase in synchronous activity from a set number of neurons [33,44]. Our study cannot elucidate the precise source of this enhanced peak response. One possibility is that the reduced interneuron recruitment following EE may reduce local inhibition and allow more excitatory input to dominate and enhance peak responses.

If enhanced pyramidal cell excitability promotes loss of cue specificity and general enhancement of peak activity, then this enhancement may represent alterations in encoding cue-related information. Related to this, we have observed disrupted sucrose cue responding following chemogenetically enhancing mPFC ensemble excitability [17]. Additionally, increased pyramidal cell activity *in vivo* may increase behavioral control over food cues, as PL hypoactivity is known to produce the opposite effect and decreases inhibitory behavioral control [45].

Distinct PL projections play different roles in reward seeking. For instance, food cue presentations largely excite and suppress PL neuronal ensembles that project to the nucleus accumbens and paraventricular nucleus of the thalamus [18]. A caveat of this fiber photometry study is that the activity of specific neuronal ensembles and projection pathways cannot be revealed. Therefore, we cannot exclude the possibility that distinct neurons were recruited with larger or more synchronous activity following EE. Future studies should utilize tools such as miniature endoscopes that provide single-cell resolution to reveal altered encoding patterns of individual PL projection neurons [46] which are newly recruited following EE and/or the ensembles which are normally recruited under standard housing conditions.

### EE-induces inhibitory underdrive in PL interneurons

Despite EE still reactivating neurons that established the ‘original’ (prior to EE) sucrose cue memory, EE attenuated the recruitment of neurons that were not cue-tagged with tdTomato. Since tdTomato expression here is Cre-dependent, these neurons that exhibit decreased recruitment may reflect neuronal populations that were activated in the prior tagging session, but did not drive sufficient Cre expression for detectable tdTomato levels.

Although we did not examine Fos in specific interneuron subtypes, one possibility here is that EE reduces the recruitment of parvalbumin (PV)-expressing interneurons. These neurons inhibit activity of nearby pyramidal cells [47] and facilitate the suppression of sucrose seeking [48]. Hence, EE-mediated decreases in the recruitment of PV-expressing interneurons, may contribute to decreased

PL Fos expression as well as persistent activation and generally elevated *in vivo* activity of pyramidal cells. Further studies are required though, to elucidate the role of PV-expressing interneurons in our observed behavioral and neuronal effects.

### Summary and future directions

Our findings deepen our understanding of PL’s behavioral role. Here we demonstrated how EE dynamically modulated PL neuronal excitability- and activity-related properties, and engaged mechanisms linked to excitatory overdrive and inhibitory underdrive. We revealed how a non-pharmacological intervention can harness prefrontal circuits that are relevant for understanding the ‘anti-food seeking’ network. Critically, EE boosted the excitability of cue-reactive neurons that ‘originally’ established a sucrose cue memory, indicating its ability to modify behaviorally relevant neurons. Since these neurons exhibited decreased rheobase, future investigations could identify sources of this decrease and investigate factors such as alterations in sodium channel conductance and subunit composition, using ‘Patch-seq’ that combines whole-cell recordings with transcriptomic analysis [49].

Furthermore, PL ensembles activated during sucrose self-administration project to reward- and motivation-relevant structures such as the BLA and nucleus accumbens^26^. Hence, an interesting future investigation is to reveal whether EE exerts its ‘anti-food seeking’ actions via modifying the downstream connectivity of cue-reactive PL ensembles in these structures. In summary, our current study sheds further light on the complex nature of excitation- and inhibition-relevant PL adaptations that might contribute to reducing the impact of food cues and thus contribute to EE’s ‘anti-craving’ effects. Such neurons may serve as potential neurophysiological targets that guide development of novel therapeutics that control food cravings.

## Funding and Disclosure

This research was supported by the UK Medical Research Council (MR/T03260X/1), BBSRC (BB/X000427/1) and the BBSRC SoCoBio Doctoral Training Programme (BB/T008768/1). Dr. Kate Peters is funded by the Leverhulme Trust. Dr. Nobuyoshi Suto is PARTIALLY SUPPORTED by U01DA055017 from the National Institute on Drug Abuse/National Institute of Health, USA.

## Competing interests

The authors have nothing to disclose.

## Data availability statement

The datasets generated during and/or analyzed during the current study are available from the corresponding author on reasonable request.

## Acknowledgments

We would like to thank Dr. Andre Chagas and Prof. Hans Crombag (University of Sussex), Drs. Gabriella Margetts-Smith (University of Bristol), Leslie Ramsey, and Alex Hoffman (NIDA/NIH) for technical support and/or discussions and Prof. James McCutcheon (Arctic University Tromsø, Norway) for providing open access analysis packages (https://github.com/mccutcheonlab/trompy). We also thank Dr. Joseph Ziminski (Sainsburys-Wellcome Center). We thank Drs Bryan Roth, Karl Deisseroth and James M Wilson for gifting their plasmids to Addgene and allowing us to purchase and use them via Addgene.

## Author Contributions

E.K. conceived and designed the research. K.Z.P., Z.P., R.A., E.C.W., and S.B.K. performed the behavioral procedures. K.Z.P., Z.P., R.A., E.C.W., S.B.K., N.S. and E.K. analyzed the behavioral results. K.Z.P., Z.P., R.A., E.C.W., S.B.K., and O.T. performed the histological procedures. K.Z.P., Z.P., E.C.W., S.B.K., O.T. and E.K. analyzed the histological results. K.Z.P., Z.P., N.S. and E.K. wrote the manuscript. K.Z.P. performed the fiber photometry procedure and analyzed the results. Z.P. performed the chemogenetic silencing procedure and analyzed the results. R.A. and O.S. performed the electrophysiology procedures. R.A., O.S., and E.K. analyzed the electrophysiology results. E.K. coordinated this work. All authors helped with data interpretation and manuscript editing.

## Notes

### Competing Interest Statement

The authors have declared no competing interest.

