## Supplemental information for "Prelimbic cortical excitatory overdrive and inhibitory underdrive accompany environmental suppression of food seeking"

### Materials and methods

#### Animals

Both male and female mice exhibited similar levels of locomotor activity (**Fig. S1a**) and similar attenuated sucrose seeking following EE (**Fig. S1b**), and our pilot experiments revealed that female mice showed reductions in Fos similar to our previous study [1]. Hence, we used both male and female mice in Experiments 1, 2 and 4 (data was collapsed) and only female mice in Experiments 3, 5, and 6. All mice were randomly allocated to their respective conditions. In Experiments 1, 5, and 6 C57BL/6J mice purchased from Charles River UK were used. In Experiments 2-4 *FosTRAP2* mice (Strain #:030323, Jackson Labs) were bred in house and for Experiments 3 and 4 double transgenic *FosTRAP2* mice were bred with *Ai14* mice (Strain #007914; Jackson Labs).

Genotypes were confirmed using the Transnetyx service (www.transnetyx.com). All mice were group housed and aged 8-12 weeks at the beginning of experiments and were housed in groups of 2-4 per cage and under a 12 h light/dark cycle (lights on at 7:00 A.M.) at a maintained temperature of 21 ± 1°C and 50 ± 5% relative humidity. One week before conditioning and until experiment completion mice were food restricted to 90% of their *ad libitum* body weight. All experiments were conducted in accordance with the UK Animals (Scientific Procedures) Act 1986 (ASPA), amended 2012, under the project license PP5152647 and received ethical approval from the University of Sussex Animal Welfare and Ethics Review Board (AWERB).

#### Magazine Training and Pavlovian Conditioning

Similar apparatus and procedures were used as previously described [2,3]. Briefly, behavioral training and testing were conducted in mouse conditioning chambers (15.9 x 14 x 12.7 cm; Med Associates, Vermont, USA) with illuminated house lights and each housed within a sound-attenuating cubicle. The chamber's front and rear access panels and ceiling were constructed from clear Plexiglas and side walls were made from removable aluminum panels atop a stainless-steel grid floor. A syringe pump dispensed 10% sucrose solution (serving as the unconditioned stimulus (US) into a recessed magazine receptacle. In fiber photometry experiments where mice were tethered during recordings, we 3D printed a modified magazine (not recessed) to allow free movement and access for mice with head implants. The conditioned stimulus (CS) was an auditory clicker created by a mechanical relay which provided a broad-frequency (0–15 kHz) sound at ~50 dB. Med-PC IV (Med Associates) software controlled all experimental parameters and data collection.

Mice first underwent a single magazine training session during which they received 40 ~15 µl sucrose solution deliveries on a random interval-30 (RI-30) schedule. Next, mice underwent 10-12 conditioning sessions, 2 times daily in the morning (8:00 A.M) and afternoon (1:00 P.M). Each acquisition session comprised of 6 presentations of the CS (120 second clicker) separated by RI-120s inter-trial intervals (ITI). During each CS period, ~15 µl deliveries of 10% sucrose solution were delivered on a RI-30 s schedule across the CS duration (average 4 US deliveries per CS trial). Number of head entries into the magazine during the CS and ITI were detected using infrared beam break sensors across the magazine and recorded in MED-PC IV software, as was time spent in seconds in the magazine.

At 3-7 d (Experiments 1, 5, and 6) and >2 weeks (Experiments 2 and 4) following the last acquisition session, mice were exposed to 24 h EE or left in standard housing (SH) **(Figs 1a-b)**. In Experiments 1, 6 and 7, mice then underwent a single test session to assess cue-evoked food seeking. Six CS's were presented under the same schedule as conditioning, but in the absence of sucrose delivery (i.e. under extinction conditions). The number of head entries into the magazine during the CS and ITI were recorded.

Timelines for Experiments 1, 5 and 6 are shown in **Figures 1a and b.** Timelines for Experiments 2, 3, 4 are indicated in **Figures 2, 3 and 4**, respectively.

**Environmental enrichment (EE)**

Standard housing (SH) consisted of a small standard cage (15.9 x 14 x 12.7 cm) with basic nesting material and a wooden chew block. Environmental enrichment (EE) housing consisted of larger cages with 3 tiers (40 x 26 x 53 cm), with connecting tunnels, a separate sleeping pod, two exercise wheels, multiple forms of nesting and bedding material, red plastic houses, cardboard tunnels, wooden chew bars and Lego bricks (**Fig. 1a**) [1]. Mice were randomly allocated to EE and SH groups and all EE exposure was for 24 h.

**Locomotor activity testing**

Locomotor testing was performed in square clear acrylic locomotor chambers (3mm thick, 20 x 20 x 20 cm) each housed in a sound attenuating cubicle. Webcams were situated directly above the chambers. Automated tracking was performed using Ethovision tracking software (Noldus, RRID:SCR_000441). The total distance traveled (m) was measured for 60 minutes. To assess possible differences in anxiety-like behavior [4] in SH and EE mice, we also measured the percentage of time spent in the central portion (defined as a 10 $cm^2$ central square area) versus the time spent at edges around the walls of the arena **(Fig. S1b)**.

**Immunohistochemistry and Fos quantification**

For Fos staining, (Experiments 4 and 5) ninety minutes following initiation of the final test session – or following EE if no test session was given (**Fig. 4h**) – mice were anaesthetized with 200 mg/kg sodium pentobarbital and transcardially perfused with phosphate buffered saline (PBS 1X; 137 mM NaCl, 10 mM $PO_4^{3-}$, 2.7 mM KCl, pH 7.4) and then 4% paraformaldehyde in PBS 1X. Brains were post-fixated overnight in 4% PFA, then cryoprotected with 30% sucrose in PBS 1X before being frozen on dry ice and stored at -80°C. Coronal sections were taken at 30 µm thickness using a Leica CM1900 cryostat. For Exp 2 and Exp 4a slices contained the medial prefrontal cortex (corresponding to approximately AP 2.13-1.73; [5]). All slices were stored in PBS 1X with 0.02% sodium azide at 4°C.

Free-floating sections were washed in PBS 1X three times for 10 min, then blocked for 60 minutes (blocking solution: PBS 1X, 0.2% Triton X-100, 3% normal goat serum (cat# S-1000, RRID:AB_2336615; Vector Laboratories). Sections were then incubated in 1/300 anti-Fos primary antibody (cat# 2250, RRID:AB_2247211; Cell Signaling Technology), and for tracing experiments 1:1000 anti-GFP (catalog #ab13970, RRID:AB_300798; Abcam, made in blocking solution). Slices were incubated overnight at RT overnight on a shaker.

The following day, slices were washed in PBS 1X three times then incubated in their corresponding secondary antibodies for 90 minutes. Secondaries were: anti-rabbit Alexa 488 (for Fos detection in Exp 6, catalog #10729174, RRID:AB_2576217; Fisher) and anti-chicken 568 (for GFP detection, catalog #SAB4600039, RRID: AB_2631230; Sigma-Aldrich). All secondaries were used at 1/300 dilution and made in PBS 1X. The fluorescent dye DAPI (catalog #D-9542, Sigma-Aldrich) was used for nuclear staining at a 5 µM concentration, slices were incubated for 45 minutes. Slices were mounted on microscope slides (catalog # 11562203; Fisher UK) air-dried, and coverslipped with Fluoromount G (catalog #00-4958, RRID:SCR_015961, Invitrogen).

Fluorescence images of Fos staining (or native fluorescence from mCherry (Exp 2), mRuby (Exp 5), and GFP staining (Exp 6) from left and right hemispheres of the PL from 1–2 coronal sections per animal, corresponding to approximately bregma 2.13 to 1.73 in Paxinos and Franklin (2001)[5], were captured using a QI click camera (Qimaging) attached to an Olympus Bx53 microscope. Fos+ nuclei were quantified using iVision software (version 4.0.15, RRID: SCR_014786; Biovision Technologies). Images of Fos immunoreactive (IR) nuclei in the PL cortex were digitally merged into a single image from 10 Z-stacks using iVision, at multiple focal planes (10 µm thickness). Representative images of the regions of interest (ROIs) for cell counts and for GCaMP6f expression visualized by its native fluorescence were taken using a QI click camera (Qimaging) attached to an Olympus Bx53 microscope running iVision software (version 4.0.15, RRID: SCR_014786; Biovision Technologies).

### *Ex vivo* electrophysiology

#### *Brain slice preparation*

One day following EE (or no EE in control SH mice), mice brains were rapidly removed and immersed into near-freezing oxygenated high-sucrose artificial cerebrospinal fluid (S-aCSF) with the following composition (mM): 60 sucrose, 87 NaCl, 2.5 KCl, 3 $MgCl_2$, 0.5 $CaCl_2$, 26 $NaHCO_3$, 1.25 $NaH_2PO_4$ $H_20$, and 10 D-glucose, saturated with 95% $O_2$ /5% $CO_2$. Coronal slices containing the prelimbic cortex (PL) of the mPFC. (250-300 µm thick sections; ~bregma 1.5-2.4 mm) were sliced using a Leica VT1200S vibrating tissue slicer. Brain slices were then stored in a holding chamber containing carbogenated-aCSF in a prewarmed water bath at ~34℃ for 30 minutes. Then cooled to room temperature for a further 10-30 minutes until the end of the recording session. For recordings, slices were transferred to a recording chamber and perfused at 2–3 ml/min with 30–32°C standard aCSF containing (mM): 125 NaCl, 2.5 KCl, 2 $CaCl_2$, 1 $MgCl_2$, 26 $NaHCO_3$, 1 $NaH_2PO_4$ $H_20$, 10 D-glucose, and bubbled with 95% $O_2$ /5% $CO_2$. Neurons were visualized with differential interference contrast using an Olympus BX51WI microscope.

#### *Whole-cell recordings*

Whole-cell recordings on PL layers V-VI pyramidal cells from PL tdTomato+ (ensemble) and tdTomato– (non-ensemble) pyramidal cells from *FosTRAP2:Ai14* mice in Experiment 3 (**Fig. 3**) were performed using borosilicate capillary glass-pipettes (1.5 mm outer diameter, 0.86 mm inner diameter), for intrinsic excitability recordings intracellular solutions consisted of (in mM): 125 K-gluconate, 10 KCl, 2 $MgCl_2$, 0.1 $CaCl_2$, 10 HEPES, 1 EGTA, 2 Mg-ATP and 0.2 Na-GTP (pH 7.2-7.4). Pyramidal cells were identified based on their morphology and/or characteristic firing properties similar to previous studies [2,3,6]. Ensemble neurons were identified based on tdTomato expression and were confirmed to be tdTomato+ during recording by colocalization of tdTomato and Alexa 488 (**Fig. 3b**).

Pipette resistances ranged from 3 to 7 MΩ. Putative pyramidal cells were identified based on their morphology and characteristic firing properties in response to current injection[6]. Data were collected with a Multiclamp 700B amplifier (Molecular Devices), A/D board (PCI 6024E; National Instruments) and WinWCP Software (courtesy of Dr. John Dempster, University of Strathclyde, Glasgow, UK. John Dempster (**http://spider.science.strath.ac.uk/sipbs/software_ses.htm**). Signals were amplified, filtered at 4 kHz and digitized at 10 kHz. The Hum Bug noise eliminator (Quest Scientific) was used to reduce noise. Neurons were visualized with differential interference contrast using an Olympus BX51WI microscope attached to a Revolution XD spinning disk confocal system (252, Andor Technology) for fluorescence microscopy.

*Intrinsic excitability recordings*

The current clamp protocol consisted of 1000 ms current injections from −200 pA to 400 pA incrementing in 10 pA steps for pyramidal cell ensemble and non-ensemble neurons.

The liquid junction potential was −13.7 mV and was not accounted for. Spike counts, spike kinetics and input resistance were analyzed with Easy Electrophysiology software (Easy Electrophysiology Ltd). Spike threshold was measured using the third differential with EasyElectrophysiology software. The action potential (AP) peak was calculated as the difference between the AP peak and AP threshold. Half-width was measured as the AP width at half-maximal spike following cubic spline interpolation to increase sampling rate by a factor of 4. Post-spike fAHPs and mAHPs were measured ~0-3 and 10-50 ms following the AP threshold, respectively, similar to [7]. Inter-spike interval (ISI) was measured as the time between the first two AP's (ISIfirst) and the last two AP's (ISIlast) of a record firing at >10 Hz. Adaptation index was measured as the $ISI_{last}$ / $ISI_{first}$ on a record with at least 6 spikes per 600 ms (10 Hz) [8]. Input resistance was calculated from the slope of the I-V curve.

### ***FosTRAP2*** **tagging**

4-hydroxytamoxifen (4-TM) was purchased from HelloBio (HB6040) and a dose of 50 mg/kg ip was injected in all tagging experiments (Experiments 4-6). We dissolved 50 mg of 4-TM in 2.5 ml of ethanol (Thermo Fisher, 16606002). This stock solution of 4-TM (20 mg/ml in ethanol) was further diluted in a 1:4 castor oil:sunflower seed oil vehicle on the day of injection resulting in 10 mg/ ml final concentration. Animals received 4-TM (50 mg/kg) intraperitoneally 3 hours after the onset of a tag session to induce robust tagging.

*Demonstration of 4-TM dependence and stimuli-induced modulation of hM4Di tagging*

To demonstrate that activity-based tagging was 4-TM dependent and modulated by exposure to appetitive stimuli in *FosTRAP2* mice, Male and female *FosTRAP2* mice were injected with AAV-hM4Di-mCherry bilaterally in PL in Experiment 2 (**Fig. 2**). Following 3-4 weeks of recovery, mice were habituated to transfer to experimental room for five consecutive days and each day they received an intraperitoneal (i.p.) injection of saline (200 µl / mouse). We allocated mice to one of three groups: 4-TM injections following a conditioning session (4-TM+ / S), 4-TM injections (4-TM+ / HC) or no injections (4-TM- / HC) in the home cage. Following a two-day quiet period, food-restricted mice in the experimental group (4-TM+/ S) underwent a single conditioning session followed by a single i.p. injection of 4-TM (50 mg/ kg) approx. 3 hours after the session onset. Mice in control groups (4-TM+/ HC, 4-TM-/ HC) received an equivalent volume of either 4-TM or saline, directly from the home cage without the conditioning session. After the 4-TM or saline treatment, all mice were kept in a quiet room for further 3 hours. Approximately 3 weeks after the tagging session, all mice were transcardially perfused, followed by brain removal and mCherry immunohistochemistry to determine hM4Di expression in PL (**Fig. 2d**).

*hM4Di-mediated silencing of PL neuronal ensembles*

Male and female *FosTRAP2* and wild-type mice underwent Pavlovian conditioning (Experiment 2, **Fig. 2c**). Mice received an i.p. injection of saline (200 μl / mouse) on the last 4 conditioning days. Five days following conditioning a group of *FosTRAP2* and a group of WT mice were injected with 4-TM following a cue-evoked sucrose seeking test to tag (or not tag in WT mice) cue-activated ensembles with hM4Di. A third group of *FosTRAP2* mice were exposed to a novel environment and were injected with 4-TM to tag neurons activated by novelty that were unrelated to sucrose cues. 4-TM was administered approx. 3 hours from the session onset. After the 4-TM injection, mice were kept in a quiet room for further 3 hours before transfer to the colony room to reduce any non-specific tagging from external stimuli. Two and a half weeks were allowed for the selective expression of the hM4Di DREADD in PL neurons activated during sucrose-seeking before cue-evoked sucrose seeking test was conducted **(Fig 2d)**. The DREADD agonist, clozapine (0.1 mg/ kg, i.p., HB1607, HelloBio) was administered 30 min before the onset of the test session to silence hM4di-expressing neurons. The dose of 0.1 mg/kg clozapine was chosen as previous studies revealed that this dose does not affect sucrose seeking [2] and Fos expression [9]. In line with past studies, in this study we demonstrated in wild-type mice that 0.1 mg/kg clozapine had no effect on cue-evoked sucrose seeking (**Fig. S2**) and thus was behaviorally inert. Therefore, the result from this control experiment demonstrates that any observed behavioral effects on sucrose seeking in our study was primarily due to the DREADD-mediated neuronal manipulation and not due to any potential confounding effects of clozapine.

*tdTomato-tagging of neurons during sucrose seeking in FosTRAP2:Ai14 mice*

We labelled neurons activated during cue-evoked sucrose seeking with tdTomato in *FosTRAP2:Ai14* mice to examine re-activation patterns of PL ensembles (Experiment 4, **Fig. 4a**) and to characterize the excitability properties of ensemble neurons (Experiment 3, **Fig. 3**). Mice underwent tagging with 4-TM 1 week following conditioning, similar to the ensemble silencing experiment (Experiment 2) and were examined approximately 2 to 2.5 weeks later for Fos expression patterns (Experiment 4) or excitability properties (Experiment 3) immediately following 1d EE or no EE (SH condition).

**Virus information**

Adeno-associated viruses:

**Experiment 2–**For expression of the inhibitory DREADD hM4Di in silencing experiments in FosTRAP2 mice we used AAV5-hSyn-DIO-hM4Di-mCherry[10] (viral titer 7 x $10^{12}$ vg/ml, Addgene #44362) diluted ¼ with sterile saline.

**Experiment 5** – For expression of mRuby in inhibitory neurons we used AAVDJ/8/2-mDlx-mRuby (viral titer 6.9 x $10^{12}$ vg/ml; catalog # v242-DJ/8, Viral Vector Facility (VVF) of the Neuroscience Center Zurich (ZNZ), constructed using elements from mDlx-HBB-chI: Addgene #83900 and mRuby3: Addgene #85146) we diluted this virus 1/10 with sterile saline.

**Experiment 6** – Genetically encoded calcium sensor AAV1-CamKII-GCaMP6f (viral titer 4.5 x $10^{12}$ vg/ml; catalog # 100834, Addgene) diluted 1/4 with sterile saline.

**Surgical procedures**

Mice (between 8 and 11 weeks old) were anaesthetized with isoflurane (4% in 1L/min oxygen for induction, 1.5-2% for maintenance), they were mounted onto a stereotaxic frame (RWD Life Science) and received pre-operative analgesia (s.c Meloxicam) as well as 200ul saline (s.c). Depilatory cream was applied to remove hair around the incision site and the scalp was cleaned with ethanol and iodine. A thermostatic blanket was used to maintain body temperature (37-38°C) throughout. An approximately 1cm incision was made with a scalpel and hole(s) was drilled above the area(s) of infusion. A sterile pulled glass pipette (Drummond PCR Micropipets 1-5ul) was attached to a syringe pump (World Precision Instruments, UMP3 micropump) and then filled with virus. Viral infusion pipette was slowly lowered to the desired DV co-ordinate and 500 nl of virus was injected at 100 nl per minute (300 nl in the case of retrograde tracing experiments). A 5-minute wait period allowed the virus to diffuse at the pipette tip before it was removed. For mRuby expression to label inhibitory cells and for DREADDs silencing experiments bilateral injections were made in the PL cortex (AP +1.8 mm, ML ± 0.35 mm, DV: -2.4 mm relative to bregma). For retrograde tracing experiments using retrograde-GFP and retrograde-RFP viruses these were injected unilaterally into the nucleus accumbens (NAc: AP +1.2 mm, ML ± 1.1 mm, DV: -3.9 mm from brain surface) and periventricular thalamus (PVT: 20° angle, AP -1.36 mm, ML ± 1.13 mm, DV: -3.3 mm relative to bregma). For viral injection only experiments the skin was sutured with polypropylene non-absorbable suture (Prolene 6-0, VetTech). For fiber photometry experiments AAV-CamKii-GCaMP6f was injected unilaterally into the PL (co-ordinates as above) and an optic fiber cannula (Doric: MFC_400/430-0.66_5mm_MF2.5_FLT) was positioned 0.1 mm above the injection site. A layer of radiopaque dental cement was used to cement the optical cannula in place (C&B Super-Bond, Prestige Dental) and a headcap was formed using dental acrylic (DuraLay, Reliance). Mice were left 3-5 weeks for recovery and viral expression before experiments began.

**Fiber photometry recordings (Experiment 6)**

Fiber photometry equipment consisted of 2 LED light sources (and their drivers) integrated into this set up 470 nm blue for excitation of GCaMP and 405 nm isosbestic (Lx405 and Lx465; Tucker Davis Technologies. These light sources were sinusoidally modulated at 210 and 330 Hz, respectively. Both light paths were directed into a 6 port filter cube (FMC6_IE(400-410)_E1(460-490)_F1(500-540)_E2(555-570)_F2(580-680)_S, Doric Lenses) and via dichroic mirrors inside directed to the mouse through a low autofluorescent optical patch cord (MFP_400/430/1100-0.57_2m_FCM_MF2.5_LAF, Doric Lenses). Bronze sleeves (info) were used to mate the patch cable to the ferrule on the mouses head for recordings. Emitted fluorescence was collected through this patch cable and measured using an integrated photoreceiver (LxPS2; Tucker Davis Technologies). The light power of 465 nm and 405 nm wavelengths was set to 20-40 uW. A signal processor (RZ10x; Tucker Davis Technologies) and Synapse software (Tucker Davis Technologies) controlled all LEDs and acquired all signals. Online demodulation was performed in Synapse to separate isosbestic and calcium-modulated responses elicited by blue or violet light sources. Demodulated signals were acquired at 1017 Hz. Behavioral events (head entries, clicker stimuli) were marked by TTL pulses generated from MED-PC hardware (SuperPort card, DIG-726TTL-G) and sent via MED-PC using a data communication cable (MedAssoc_CBL, Tucker Davis Technologies). The processor was controlled by a WS4 PC (Tucker Davis Technologies) and all data files were stored locally and backed up to external hard drives. Before recordings optical patch cables were photobleached at each wavelength (405 nm and 465 nm) for 4 hours. Before experiments began, mice were habituated to restraint and patch cord connection (10 mins per day for 3 days).

**Data Analysis**

All statistical tests were performed using Prism software (RRID:SCR_002798; GraphPad 10 Software). For all experiments, data was analyzed using ANOVAs, followed by post-hoc testing or pair-wise comparisons (see below for details). Group data are presented as mean ± SEM. Data points exceeding +/-2 SDs from the mean were excluded as outliers.

*Experiment 1: Cue-evoked sucrose seeking and open field activity*

Total head entries into the sucrose-delivery magazine with data from males and females combined during CS and ITI presentation during acquisition and extinction were analyzed using a two-way ANOVA including the within-subjects factors of CS (CS, ITI) and session. Sucrose seeking test data following EE were analyzed using a two-way ANOVA using within-subjects factor of CS Presentation (CS, ITI) and between-subjects factor of EE (EE, SH). Open field data was analyzed using a Mann-Whitney U test.

*Experiment 2: Demonstrating 4-TM dependency of tagging and hM4Di-mediated silencing of ensemble neurons.* A 1-way ANOVA was used to assess differences in hM4Di-mCherry expression across mice in different groups (4-TM+ / S; 4-TM+ / HC; and 4-TM- / HC). To analyze the subthreshold effects of clozapine on cue-evoked sucrose seeking, a 2-way ANOVA with within-subjects factor of CS and between-subjects factor of Clozapine was used and for demonstrating effects of clozapine on ensemble silencing, a 2-way ANOVA was used with within-subjects factor of factor of CS and between-subjects factor of Group.

*Experiment 3: Excitability of PL ensemble neurons.* Firing capacity of ensemble neurons were analyzed using a 3-way ANOVA with between-subjects factors of EE (EE, SH), tdTomato (tdTomato+ vs –) and within-subjects factor of Current. Active and passive membrane kinetics were analyzed using a two-way ANOVA with factors of EE (EE, SH) and tdTomato (tdTomato+ vs –).

*Experiment 4: Examining re-activation patterns of tdTomato-tagged sucrose seeking ensembles in* FosTRAP2:Ai14 *mice and baseline Fos.* A 2-way ANOVA with factors of within-subjects factor of CS and between-subjects factor of EE was used to analyze effects of assignment to EE and SH groups in Cue test 1 and to determine the effects of EE on sucrose seeking in Cue test 2. Cell counts for tdTomato+ and/or Fos+ cells were analyzed using an unpaired, 2-tailed t-test between housing conditions (EE vs SH).

*Experiment 5: EE effects on cue-evoked Fos in interneurons.* An unpaired, 2-tailed *t* test and Mann-Whitney U test was conducted to assess differences in Fos+ and/or mRuby+ cell counts per square millimeter and percentage of Fos+ cells expressing mRuby, respectively, between housing conditions (EE vs SH).

*Experiment 6: Fiber photometry recordings during conditioning and test.* Fiber photometry data (fluorescent signals and behavioral timestamp events) was extracted using Python packages from TuckerDavisTechnologies (TDT: https://pypi.org/project/tdt). Pre-processing, baseline correction and z-score normalization was carried out using the Trompy package (Prof. James McCutcheon: https://github.com/mccutcheonlab/trompy). A peak detection algorithm was written in Python based on previous studies [11,12]. Signals were filtered to remove high amplitude events (local maxima two median absolute deviations (MAD) above the median of a 10 second moving median). Peaks / transients were detected as local maxima more than 3 MADs over the median. Amplitude and frequency were compared across ITI and CS periods in SH and EE mice using 2-way repeated measures or mixed ANOVA with the within-subjects factor of CS (and session) and between-subjects factor of EE. Head entry aligned averages were calculated and signal plots were made using custom written Python scripts adapted from several functions within the Trompy Python package. For each mouse, activity traces during head entries were aligned to head entry onset and 2 seconds following then averaged. To quantify activity peaks for head entry aligned activity we measured the maximum z-score value within the head entry window (2 seconds following entry). We compared the average amplitude of head entry related activity for each mouse during ITI vs CS.

**Supplemental Figures**

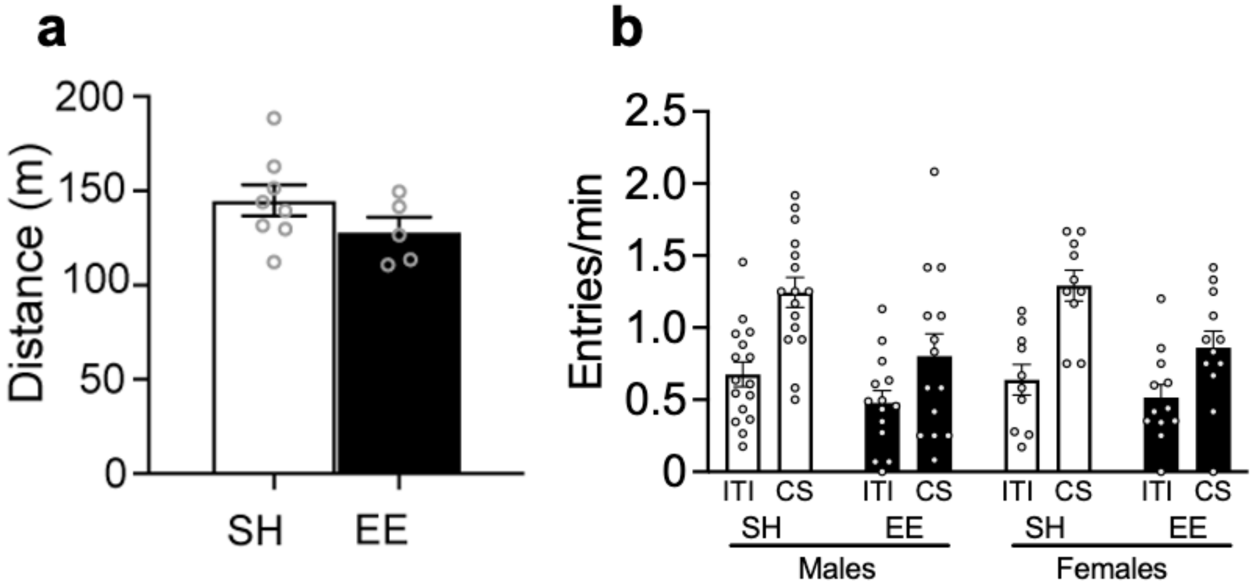


**Figure S1**.

**a** Lack of locomotor activity effects following EE (n=6-8/group). **b** Lack of sex differences on EE effects during cue-evoked sucrose seeking (Sex X CS X EE: $F_{1,18}$ = 0.08810, p=0.7700) but EE modulates cue-evoked sucrose seeking (CS X EE: $F_{1,18}$ = 6.380, p<0.05; n=6-8/group). All data are expressed as mean ± SEM.

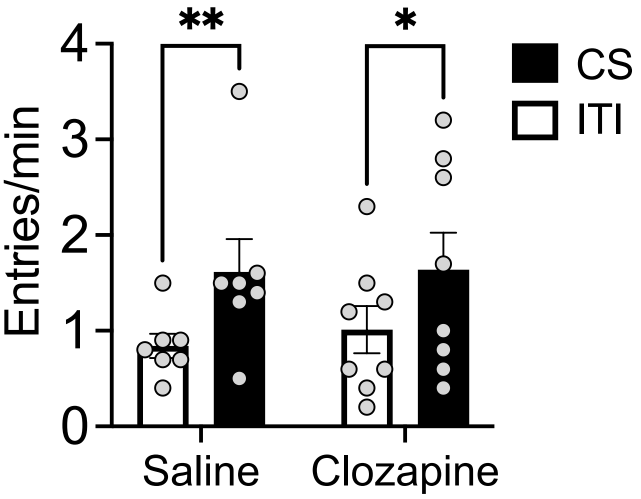


**Figure S2**

Lack of clozapine effects on cue-evoked sucrose seeking (n=7-8/group). Clozapine (0.1 mg/kg, i.p.) did not modulate cue-evoked sucrose seeking as revealed by lack of Clozapine x CS interaction (**$F_{1,13}$** = 0.226, p=0.643) while revealing a main effect of CS ($F_{1,13}$ = 20.53, p<0.001). *p<0.05 and **p<0.01. All data are expressed as mean ± SEM.

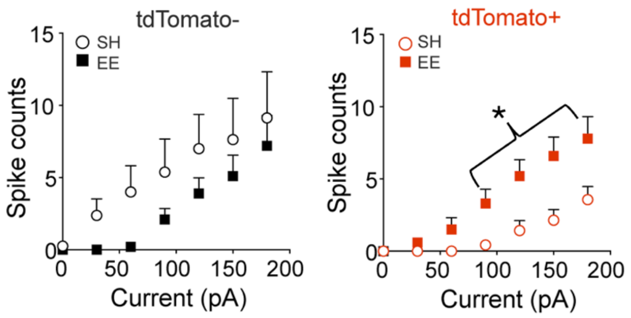


### Figure S3

The firing capacity of tdTomato+ and tdTomato– cells across EE and SH conditions. (EE: n=10/4, tdTomato+ and 9/5, tdTomato–; SH: n=7/4, tdTomato+ and n=8/3, tdTomato–). *n* = total number of cells/total number of mice. $^{*}p<0.05$; EE vs SH. All data are expressed as mean ± SEM.

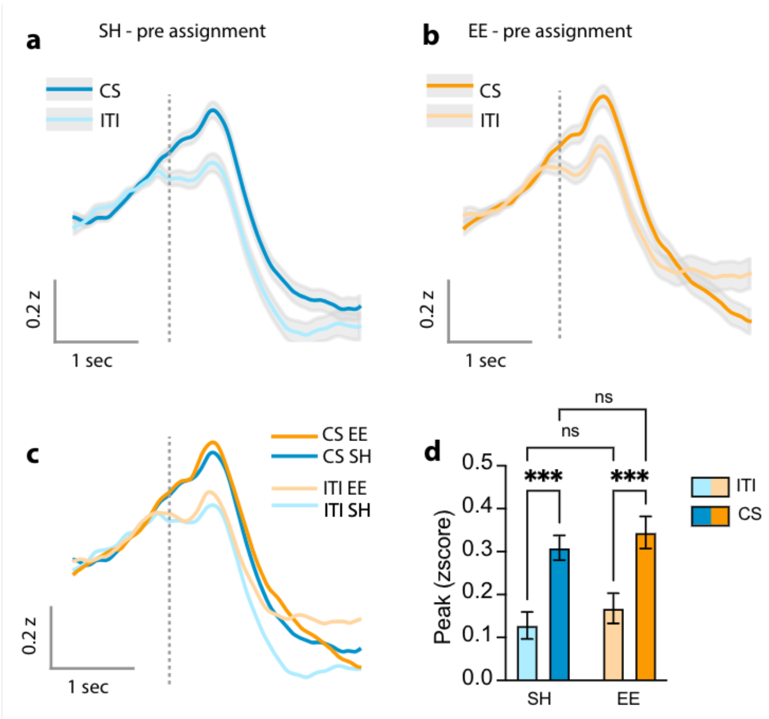


### Figure S4

Mice assigned to the EE and SH conditions show similar cue-evoked increases in PL activity *in vivo.*

**a** - Head entry aligned activity for CS and ITI in mice which were assigned to the standard housed condition on final conditioning day (shaded error +/- SEM)

**b** - Head entry aligned activity for CS and ITI in mice which were assigned to the environmentally enriched housing condition (shaded error +/- SEM)

**c** - Average activity traces for each group and stimulus compared

**d** - Average peak for head entries during ITI or CS in SH and EE mice before group assignment. There was a significant effect of stimulus (ITI vs CS) with responses to CS significantly higher than ITI in both groups (***p<0.001). There were no significant effects of housing condition or interaction. Error bars represent mean+/- SEM.

**Table S1:** Basic membrane properties from PL ensemble and non-ensemble neurons following EE.

| | tdTomato– SH vs EE Mean+/- SEM | p Value | tdTomato+ SH vs EE Mean+/- SEM | p Value |
|---|---|---|---|---|
| Break In Resting Vm (mV) | -59.80 ± 2.97 vs -62.89 ± 3.22 | 0.4738 | -63.42 ± 2.12 vs -65.09 ± 3.13 | 0.7076 |
| Rheobase (pA) | 61.3 ± 27.2 vs 102.0 ± 19.4 | 0.1522 | 137.1 ± 18.6 vs 77.0 ± 13.4 #* | 0.0454 |
| *R*i (mΩ) | 159.4 ± 17.9 vs 133.3 ± 6.51 | 0.2869 | 124.8 ± 13.06 vs 153.0 ± 23.9 | 0.2697 |
| AP peak (mV) | 37.07 ± 1.71 vs 25.31 ± 2.81 †* | 0.0131 | 27.41 ± 1.71 vs 24.57 ± 3.92 | 0.5458 |
| Latency to First AP (s) | 0.43 ± 0.04 vs 0.44 ± 0.03 | 0.7199 | 0.40 ± 0.02 vs 0.45 ± 0.03 | 0.2725 |
| AP Threshold (mV) | -31.37 ± 8.28 vs -33.03 ± 2.88 | 0.6909 | -34.36 ± 1.07 vs -27.81 ± 6.11 | 0.5048 |
| AP Half-width (ms) | 1.49 ± 0.25 vs 1.03 ± 0.06 †* | 0.0195 | 1.02 ± 0.05 vs 0.89 ± 0.10 | 0.5345 |
| fAHP (mV) | -5.62 ± 6.02 vs -10.79 ± 1.18 | 0.2621 | -9.06 ± 1.40 vs -17.06 ± 6.22 | 0.6457 |
| mAHP (mV) | -24.51 ± 6.64 vs -18.51 ± 0.53 | 0.4017 | -18.74 ± 0.73 vs -18.94 ± 2.30 | 0.2576 |
| First ISI (s) | 0.06 ± 0.004 vs 0.07 ± 0.01 | 0.4284 | 0.04 ± 0.01 vs 0.04 ± 0.01 | 0.9563 |
| Final ISI (s) | 0.11 ± 0.01 vs 0.11 ± 0.02 | 0.6755 | 0.15 ± 0.03 vs 0.11 ± 0.01 | 0.1821 |
| Adaptation Index | 1.89 ± 0.18 vs 1.09 ± 0.21 | 0.5271 | 2.01 ± 0.72 vs 2.83 ± 1.45 | 0.5609 |
| Mean ISI (ms) | 0.10 ± 0.01 vs 0.27 ± 0.07 | 0.1143 | 0.27 ± 0.07 vs 0.35 ± 0.08 | 0.2187 |

**Table S1 Legend**

Basic membrane properties from PL ensemble (tdTomato+) and non-ensemble (tdTomato–) neurons following EE or no EE (SH) exposure. All data are expressed as mean ± SEM (#Significant interaction of EE × tdTomato; †significant main effect of EE; *significant differences between SH vs EE conditions within tdTomato+ or tdTomato– cell populations, following post-hoc Fisher's LSD test); $V_m$, Resting membrane potential; $R_i$, input resistance; fAHP, fast AHP; mAHP, medium AHP.
